# Connectional Asymmetry of the Inferior Parietal Lobule Shapes Hemispheric Specialization in Humans, Chimpanzees, and Rhesus Macaques

**DOI:** 10.1101/2021.01.26.428189

**Authors:** Luqi Cheng, Yuanchao Zhang, Gang Li, Jiaojian Wang, Chet C. Sherwood, Gaolang Gong, Linzhong Fan, Tianzi Jiang

## Abstract

The inferior parietal lobule (IPL) is one of the most expanded cortical regions in humans relative to other primates. It is also among the most structurally and functionally asymmetric regions in the human cerebral cortex. Whether the structural and connectional asymmetries of IPL subdivisions differ across primate species and how this relates to functional asymmetries remain unclear. We identified IPL subregions that exhibited positive allometric in both hemispheres, scaling across rhesus macaque monkeys, chimpanzees, and humans. The patterns of IPL subregions asymmetry were similar in chimpanzees and humans, but no IPL asymmetries were evident in macaques. Among the comparative sample of primates, humans showed the most widespread asymmetric connections in the frontal, parietal, and temporal cortices, constituting leftward asymmetric networks that may provide an anatomical basis for language and tool use. Unique human asymmetric connectivity between the IPL and primary motor cortex might be related to handedness. These findings suggest that structural and connectional asymmetries may underlie hemispheric specialization of the human brain.

## Introduction

The association cortex has expanded greatly in size and exhibits modified connectivity patterns in human brain evolution (Orban *et al*., 2006; Mars *et al*., 2017; Ardesch *et al*., 2019; Van Essen *et al*., 2019). Compared with the primary sensory and motor cortical regions, the association cortex displays disproportionate expansion in conjunction with overall neocortical volume enlargement across primates (Chaplin *et al*., 2013). Accordingly, association areas comprise a large percentage of the neocortex in human brains (Orban *et al*., 2006; Van Essen and Dierker, 2007; Donahue *et al*., 2018). Functional and neuroanatomical asymmetries are also pronounced in the human brain, appearing to be more extreme compared with other primate species, especially in the association cortex (Chance and Crow, 2007). Nevertheless, cerebral asymmetry exists not only in humans but also in nonhuman primates (Gómez-Robles *et al*., 2013; Hopkins, 2013). For example, olive baboons and chimpanzees showed population-level leftward volumetric asymmetry in the planum temporale, which is thought to be homologous to part of Wernicke’s area in humans and may have played a facilitating role in the evolution of spoken language (Spocter *et al*., 2010; Marie *et al*., 2018). Comparative studies on brain asymmetry are crucial for understanding the evolution and function of the modern human brain.

Language and complex tool use, which show considerable lateralization in the human brain, are considered to be universal features of humans (Johnson-Frey *et al*., 2005; Lewis, 2006; Binder *et al*., 2009). These specialized functions all involve the inferior parietal lobule (IPL), an area of the association cortex that represents a zone of topographical convergence in the brain (Johnson-Frey, 2004; Binder *et al*., 2009). Moreover, the IPL is one of the most expanded regions in humans compared with nonhuman primates (Orban *et al*., 2006; Van Essen and Dierker, 2007; Kaas, 2012; Ardesch *et al*., 2019). The functional diversity and expansion of the IPL imply that it contains subdivisions that may have been elaborated or developed in the ancestors of modern humans, allowing new abilities such as extensive tool use and communication using gestures (Kaas, 2012). However, due to the scarcity of data, different criteria, and methodological limitations for defining regions or subregions (Mars *et al*., 2017), whether the internal organization of the IPL differs across species and how this relates to different asymmetric functions remain unclear.

A major challenge for neuroscience is to translate results obtained using one method and in one species to other methods and other species. Although the IPL has been subdivided into distinct subregions using cytoarchitecture and this technique has provided invaluable information, cellular microstructure alone is insufficient to completely represent brain organization, especially long-range connections, which are the major determinant of regional specialization (Passingham *et al*., 2002; Caspers *et al*., 2006). Furthermore, histological methods with postmortem brains cannot be readily scaled to large populations. Recent advances in diffusion magnetic resonance imaging (MRI), which allow the quantitative mapping of whole-brain neural connectivity *in vivo*, provide an alternative technique called connectivity-based parcellation to subdivide specific regions of the brain or even the entire cortex (Fan *et al*., 2016; Eickhoff *et al*., 2018). In previous studies, this technique was successfully used to characterize IPL subdivisions in different species as well as to perform cross-species comparisons (Mars *et al*., 2011; Wang *et al*., 2020).

Previous studies have assessed asymmetries of the IPL using local characteristics, such as cortical volume, thickness, and surface area (Croxson *et al*., 2018; Kong *et al*., 2018). However, although such regional asymmetries have been identified, additional analyses are needed to address the architecture of neural connectivity (Ocklenburg *et al*., 2016). A recent “connectomic hypothesis for the hominization of the brain” suggests neural network organization as an intermediate anatomical and functional phenotype between the genome and cognitive capacities, which are extensively modified in the human brain (Changeux *et al*., 2020). The functions and interactions of brain regions are determined by their anatomical connections (Passingham *et al*., 2002). Therefore, identifying connectional asymmetries may provide new insights into the structural and functional specializations of the human brain.

This study investigated asymmetries of IPL subregions in terms of both structure and anatomical connectivity in rhesus macaques, chimpanzees, and humans. We first used connectivity-based parcellations to subdivide the IPL to reveal consistent cross-species topographical organization. We then investigated the volumetric allometric scaling and asymmetries of the IPL subregions across species. Using vertex-, region of interest (ROI)-, and tract-wise analyses, we examined asymmetries of the IPL subregions in terms of their connectivity profiles and subcortical white matter pathways to identify evolutionary changes.

## Results

### Connectivity-based parcellation

For each species, a data-driven connectivity-based parcellation was applied to group the vertices in the IPL into functionally distinct clusters based on anatomical connectivity (**Figure 1**). Because spectral clustering does not require a specific number of clusters, we iterated the number of subregions from two to twelve to search for the optimal number of subregions. To accomplish this, we identified the optimal number of subregions of the IPL by choosing the maximum number of subregions that showed a coherent topological organization across all species while balancing that by the minimum number of subregions that could be identified based on their cytoarchitectural definitions in macaques, chimpanzees, and humans (Pandya and Seltzer, 1982; Reyes, 2017). The two- to five-cluster solutions are shown in **Supplementary Figure 1**. The two- to four-cluster solutions showed a consistent rostral-caudal pattern in all three species, but in the five-cluster solution a ventral cluster emerged in chimpanzees and a dorsal cluster emerged in humans. The four-cluster solution revealed a rostral-caudal topological pattern that was consistent with previous parcellations based on cytoarchitecture and anatomical connectivity (Pandya and Seltzer, 1982; Caspers *et al*., 2006; Mars *et al*., 2011; Fan *et al*., 2016). Also, the cytoarchitectural definition of macaques revealed four subregions in the IPL (Pandya and Seltzer, 1982), which was fewer than the seven cytoarchitectural subregions of the human IPL (Caspers *et al*., 2006). Although the four-cluster solution was not the finest, especially in humans, it contained potentially valuable information about the differences between species. Furthermore, the aim of our research was not to find the “best” cluster solution for the IPL but to identify an appropriate parcellation that could shed light on the lateralization of the structure and connectivity of the IPL and its subregions in this particular sample of three primate species. As such, we chose four clusters as the optimal solution for the cross-species comparison.

**Figure 1.**
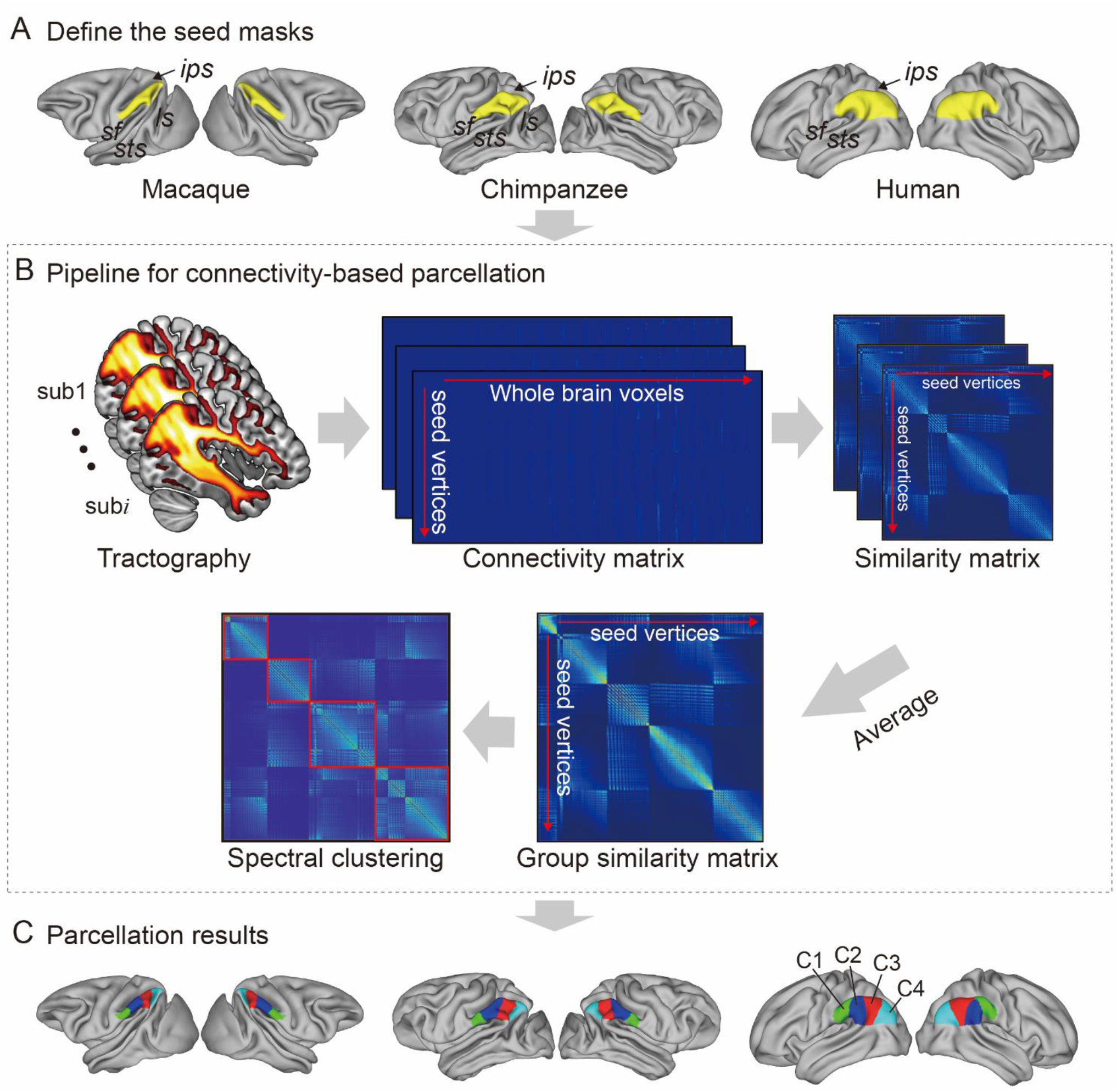
Framework of the connectivity-based brain parcellation for macaques, chimpanzees, and humans. (**A**) Defining the seed masks of the inferior parietal lobule (IPL) in surface space according to the gyri and sulci. (**B**) Connectivity-based parcellation using anatomical connectivity. Probabilistic tractography was applied by sampling 5000 streamlines at each vertex within the seed mask. Whole-brain connectivity profiles were used to generate a connectivity matrix with each row representing the connectivity profile of each seed vertex. Next, a correlation matrix was calculated as a measure of similarity between the seed vertices. Then, a group similarity matrix was calculated by averaging the correlation matrix across subjects and spectral clustering was applied to it. (**C**) Parcellation results of the IPL across species. The entire framework was applied independently for each hemisphere and each species.

It is widely accepted that the IPL contains two major cytoarchitectural divisions across species, the anterior (PF) and posterior (PG) areas (von Economo and Koskinas, 1925; Von Bonin, 1947; Bailey *et al*., 1950). Our results were consistent with this two-way parcellation and refined it into four subdivisions, specifically, two anterior clusters (the C1 and C2) in the PF and two posterior clusters (the C3 and C4) in the PG. In macaques and chimpanzees, the IPL was previously parcellated into four distinct areas based on histology (Pandya and Seltzer, 1982; Reyes, 2017) in keeping with our four-cluster solution. In humans, the IPL was cytoarchitecturally parcellated into seven distinct areas. Although we proposed a four-cluster solution that has fewer areas than the cytoarchitectural map, it is also consistent with it (Caspers *et al*., 2006). Specifically, the rostral anterior cluster (C1) is similar to the PFt and part of the PFop area defined using cytoarchitecture by Caspers et al. (2006), the caudal anterior cluster (C2) corresponds to the PF and PFm areas, the rostral posterior cluster (C3) is similar to the PGa area, and the caudal posterior cluster (C4) is similar to the PGp area. Our results did not include the PFcm area because it is located deep in the parietal operculum. Given the limited descriptions of subdivisions and connectivity of the IPL in chimpanzees, our parcellation of the IPL can depict the subregions and connectivity of the IPL in chimpanzees from an evolutionary perspective. To assess which hemisphere was dominant with respect to a given function of the human IPL subregions, we decoded the functions of the human IPL subregions from the Neurosynth database (Yarkoni *et al*., 2011) and calculated differences in the correlation values between the left and right corresponding subregions (**Supplementary Figure 2**). The term *tool* showed a much higher correlation with the left C1 than with the right C1, suggesting that the left C1 is more involved in tool use. Terms such as *tool* and *semantics* showed relatively high correlations with the left C2, whereas terms such as *nogo* and *inhibition* showed relatively high correlations with the right C2, suggesting that the left C2 is more involved in tool use and language whereas the right C2 is more involved in executive function. Terms such as *retrieval, episodic, recollection, memories*, and *coherent* showed relatively high correlations with the left C3, whereas terms such as *nogo, inhibition*, and *beliefs* were correlated with the right C3, suggesting that the left C3 is more involved in memory and language whereas the right C3 is more involved in executive and social cognitive functions. Terms such as *episodic* and *coherent* showed relatively high correlations with the left C4, whereas terms such as *spatial, attention, mentalizing*, and *relevance* showed relatively high correlations with the right C4, suggesting that the left C4 could be more involved in memory and language whereas the right C4 could be more involved in spatial attention and social functions.

### Allometric scaling and structural asymmetry of IPL subregions

When examining the relationship of the volume of each of the IPL subregions scaled against the total grey matter volume, the scaling of all the IPL subregions showed positive allometry (all slopes > 1) (**Figure 2A**). A statistical analysis revealed no significant differences between the slopes of each pair of the bilateral IPL subregions. The asymmetry indices (AIs) for the IPL subregions were calculated and are shown in **Figure 2B**. The macaques showed no significant asymmetry after Bonferroni correction for any of the subregions. The chimpanzees and humans both displayed leftward asymmetry in the rostral IPL (the C1 and C2, all *p* < .001) and rightward asymmetry in the caudal IPL (the C3 and C4, all *p* < .001).

**Figure 2.**
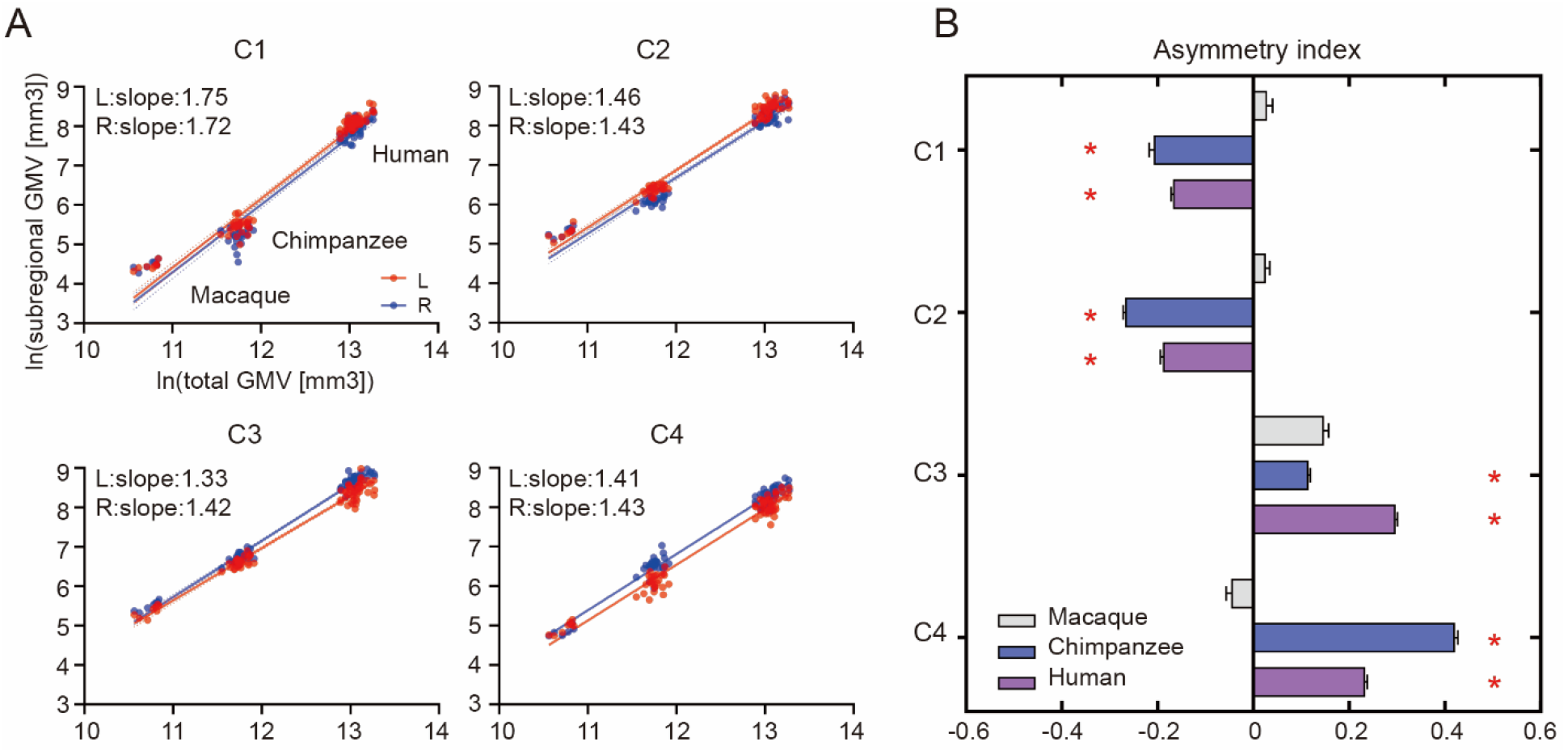
Structural allometric scaling and asymmetries of the inferior parietal lobule (IPL) subregions across species. (**A**) Volumes of the IPL subregions plotted against total cortical gray matter volume (GMV). Solid lines represent the best fit using mean macaque, chimpanzee, and human data points; dotted lines represent 95% confidence intervals. (**B**) Volumetric asymmetries of the IPL subregions. Negative asymmetry index indicates leftward asymmetry and positive index indicates rightward asymmetry. * denotes significance at the Bonferroni corrected level of *p* < .05. The error bars indicate the standard error of the mean.

### Connectional asymmetries of IPL subregions

To investigate the connectional asymmetries of the IPL subregions, we first calculated the connectivity profiles of the left and right subregions in macaques, chimpanzees, and humans using probabilistic tracking (**Supplementary Figure 3**). Visualization of the connectivity patterns of the IPL did not show significant interhemispheric asymmetry in macaque monkeys or chimpanzees but did in humans, especially in connections with the inferior frontal gyrus (IFG) and lateral temporal cortex. A vertex-wise analysis was then performed to examine the connectional asymmetry of each subregion for each species by calculating the AIs between its connectivity profiles for the two hemispheres (**Figure 3**). Additionally, ROI- and tract-wise analyses were used to examine the asymmetry of the cortical regions and subcortical white matter pathways connected to the subregions, respectively (**Figure 4;** connectivity values shown in **Supplementary Figure 4, 5**). No significant asymmetries were found in macaques in any of the statistical analyses after correction for multiple comparisons.

**Figure 3.**
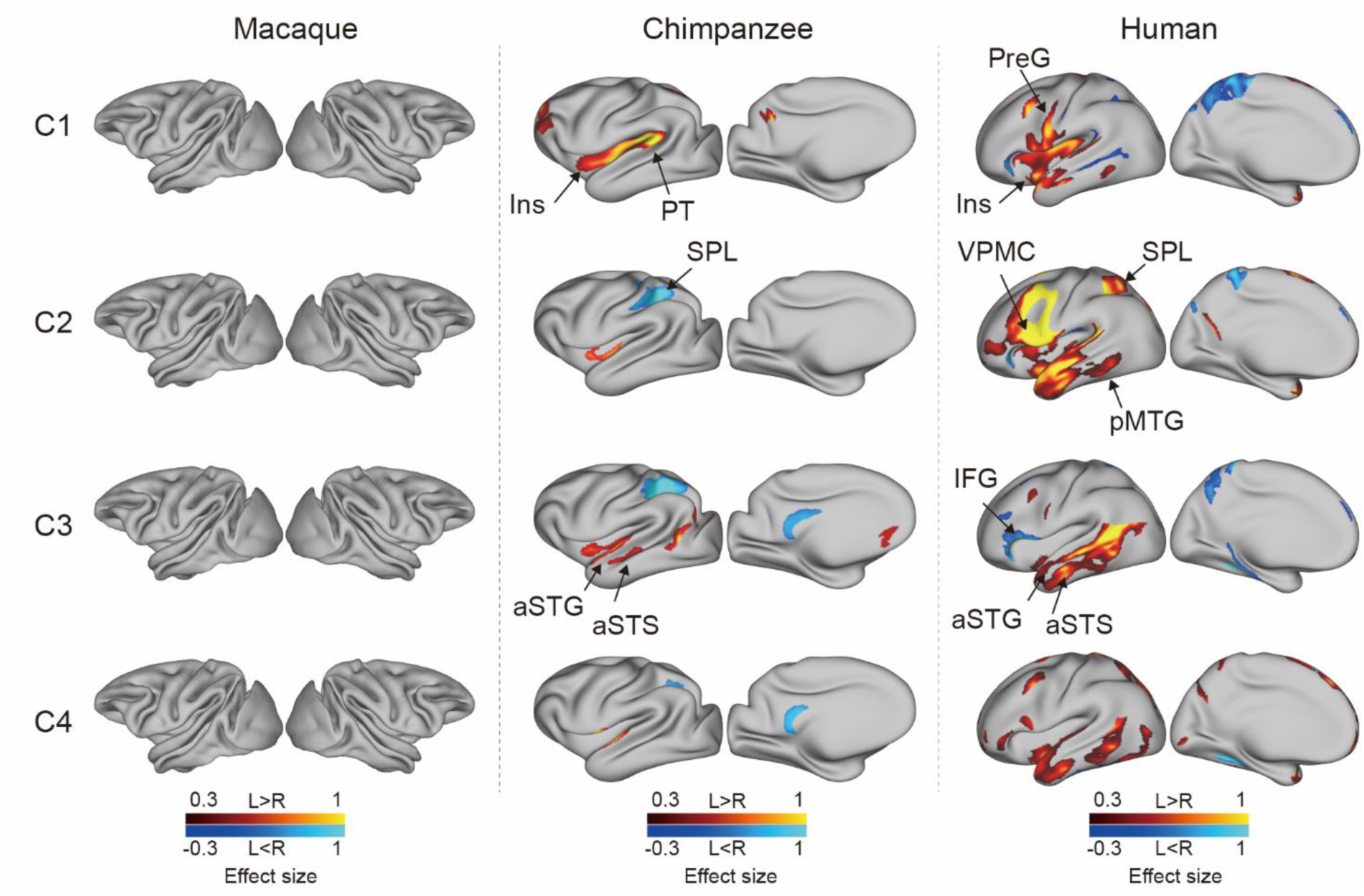
Connectional asymmetries of the IPL subdivisions in the vertex-wise analyses across species. Effect size (Cohen’s *d)* related to asymmetric connections of IPL subdivisions displayed on the left hemisphere of a species-specific standard brain (leftward asymmetry: yellow, rightward asymmetry: blue) for each species for areas showing significance at the *p* < .05 level corrected for multiple comparisons using false discovery rate correction. PreG, precentral gyrus; SPL, superior parietal lobule; aSTG, anterior superior temporal gyrus; aSTS, anterior superior temporal sulci; PT, planum temporale; VPMC, ventral premotor cortex; pMTG, posterior middle temporal gyrus; IFG, inferior frontal gyrus; Ins, insula.

**Figure 4.**
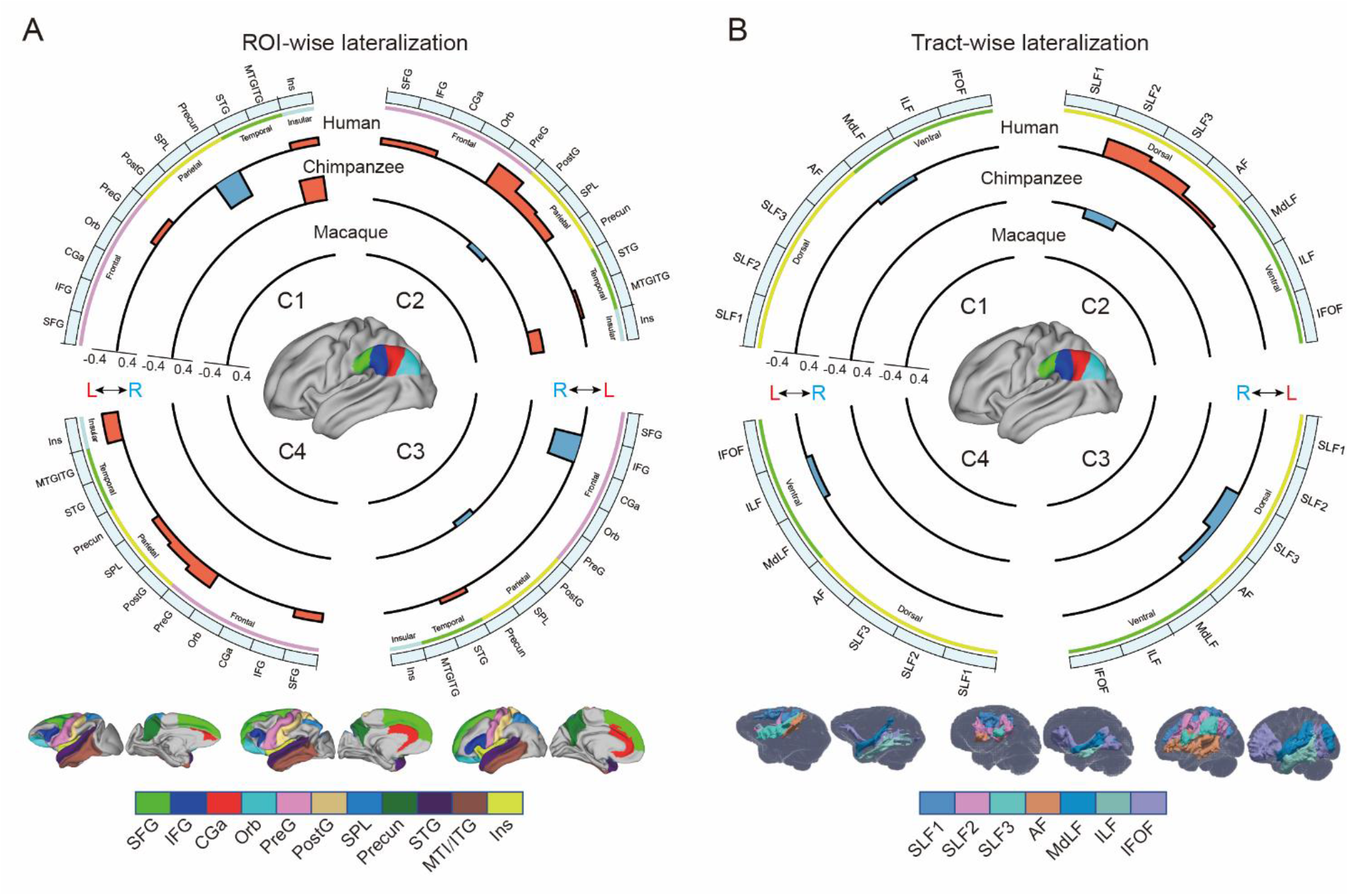
(**A**) Connectional asymmetries of IPL subdivisions in the region of interest (ROI)-wise analyses across species. Connectional asymmetry was calculated for the connections between each IPL subregion and eleven ROIs. (**B**) Connectional asymmetries of the IPL subdivisions in the tract-wise analysis across species. Connectional asymmetry was calculated for the connections between each IPL subregion and the seven tracts. For all plots, the four quadrants of each circle correspond to the four IPL subregions. The outermost circles represent ROIs or tracts. The three inner circles from inside to outside represent macaques, chimpanzees, and humans, respectively. For all plots, only the connectivity showing a significance at a Bonferroni corrected level of *p* < .05 are displayed. SFG, superior frontal gyrus; IFG, inferior frontal gyrus; CGa, anterior cingulate gyrus; Orb, orbitofrontal cortex; PreG, precentral gyrus; PostG, postcentral gyrus; SPL, superior parietal lobule; STG, superior temporal gyrus; MTG/ITG, middle temporal gyrus and inferior temporal gyrus; Ins, insula; SLF1, SLF2, SLF3, the three branches of the superior longitudinal fasciculus; AF, arcuate fasciculus; MdLF, middle longitudinal fasciculus; ILF, inferior longitudinal fasciculus; IFOF, inferior fronto-occipital fasciculus.

In chimpanzees, the C1 showed significant leftward asymmetry mainly in connections with the anterior middle frontal gyrus (MFG), anterior IFG, planum temporale, and insula. The C2 showed significant leftward asymmetric connections with the insula and rightward asymmetric connections with the superior parietal lobule (SPL) and superior longitudinal fasciculus 2 (SLF2). The C3 showed significant leftward asymmetric connections with the anterior superior temporal gyrus (STG), anterior superior temporal sulcus (aSTS), and occipitotemporal area and rightward asymmetric connections with the SPL and posterior cingulate gyrus (PCC). The C4 showed significant leftward asymmetric connections with the anterior STG (aSTG) and rightward asymmetry with the SPL and PCC.

In humans, the C1 showed significant leftward asymmetric connections with the ventral premotor and motor cortices and insula, which was consistent with regional leftward asymmetric connections with the precentral gyrus (PreG) and insula. The C1 also showed significant leftward asymmetric connections with the posterior MFG, aSTG, and posterior middle temporal gyrus (MTG) and rightward asymmetric connections with the orbital part of the IFG, posterior STS, and dorsal precuneus. The C2 showed significant leftward asymmetric connections with the posterior MFG, ventral premotor and motor cortices, SPL, anterior temporal lobe, and posterior MTG, which was consistent with regional leftward asymmetric connections with the IFG, PreG, postcentral gyrus (PostG), SPL, and STG and was supported by leftward asymmetric subcortical connections with the SLF2, SLF3, and arcuate fasciculus (AF). The C2 also showed rightward asymmetric connections with the orbital part of the IFG and posterior cingulate sulcus. The C3 showed significant leftward asymmetry mainly in the connections with the anterior IFG, SPL, and almost all the lateral temporal cortex, which was consistent with regional leftward asymmetric connections with the MTG and inferior temporal gyrus (MTG/ITG). The C3 also showed rightward asymmetric connections with the IFG, which was supported by leftward asymmetric subcortical connections with the SLF3. The C4 showed significant leftward asymmetry mainly in the connections with the IFG and anterior and posterior temporal cortex. The C4 also showed significant regional leftward asymmetric connections with the PreG, PostG, and SPL in the ROI-wise analysis.

## Discussion

In the present study, we investigated asymmetries of the IPL in the structure and connectivity of rhesus macaques, chimpanzees, and humans. In the structural analysis, the IPL and its subregions exhibited a similar pattern of positive allometric scaling between hemispheres. In addition, the chimpanzees and humans shared similar asymmetric patterns in the IPL subregions, i.e., left asymmetry in the anterior part and right asymmetry in the posterior part, whereas macaques did not display asymmetry. In the connectivity analysis, the chimpanzees showed some connectional asymmetric regions including the SPL, insula, planum temporale, aSTG, and aSTS. The humans showed widespread connectional asymmetric regions including the primary motor and premotor cortices, SPL, insula, and the entire lateral temporal lobe. These regions are associated with language, tool use, and handedness, suggesting a potential relationship between the connectional asymmetry and the functional hemispheric specialization of the human brain.

### Positive allometric scaling and structural asymmetry of IPL subregions

Brain allometry describes the quantitative scaling relationship between changes in the size of one structure relative another structure, often the whole brain or cerebral cortex (Mars *et al*., 2017; Smaers *et al*., 2017). Previous allometric studies suggested that the association cortex (prefrontal, temporal, and parietal regions) scales with positive allometry (i.e., increases in size disproportionally, or more rapidly) across primates (Passingham and Smaers, 2014; Mars *et al*., 2017). Utilizing parcellation-based delineations, a recent study provided evidence that human brains have a greater proportion of prefrontal cortex gray matter volume than other primates (Donahue *et al*., 2018) and other studies demonstrate that human prefrontal expansion is greater than would be expected from allometric scaling in nonhuman primates (Passingham and Smaers, 2014; Smaers *et al*., 2017), although some conflicting analyses remain (Gabi *et al*., 2016). In the present study, we used macro-anatomical boundaries to identify the boundaries of the IPL and a connectivity-based parcellation approach to subdivide the IPL, which helped to reveal its internal organization. We found that the bilateral IPL subregions exhibited consistent, positive allometric scaling, which suggests that allometric scaling of the internal organization of the IPL was similar and was also consistent between homotopic regions during the evolution of the IPL in anthropoid primates. With only three species in the sample, our dataset does not allow us to use phylogenetic comparative statistical methods or determine whether human IPL subregions fall significantly above allometric expectations from nonhuman primates; future research that incorporates a broad phylogenetic sample of diverse primate brains would be necessary.

We found that chimpanzees and humans showed a similar dichotomous asymmetric pattern in their IPL subregions, i.e., leftward asymmetry in the anterior portion (the C1 and C2) and rightward asymmetry in the posterior portion (the C3 and C4), but macaques did not show any asymmetry. The result in humans is consistent with a recent study using data from a large consortium showing leftward asymmetry in the supramarginal gyrus and rightward asymmetry in the angular gyrus in terms of surface area (Kong *et al*., 2018). The divergent volumetric asymmetries suggest functional heterogeneities of the IPL and emphasize the importance of analyzing subregions within the IPL. The shared asymmetric pattern also suggests that divergences in the internal organization of the IPL evolved prior to the common ancestor of chimpanzees and humans and after the common ancestor of the three species.

### Connectional asymmetries underlying human language and complex tool use

Recent neuroimaging studies have highlighted specific brain regions and pathways that may be necessary for tool use (Lewis, 2006; Stout and Chaminade, 2012). We found that humans showed leftward asymmetric connectivity between the IPL (the C2) and the primary motor cortex, ventral premotor cortex, SPL, and posterior MTG, all of which were activated in tasks related to tool use and might constitute a cortical network underlying complex tool use (Lewis, 2006). In addition, portions of this network appeared to represent part of a system that is tightly linked with language systems. The interaction between the tool use system and the language system, though with a clear left hemisphere bias, is responsible for representing semantic knowledge about familiar tools and their uses and for acquiring the skills necessary to perform these actions (Johnson-Frey, 2004; Lewis, 2006; Stout and Chaminade, 2012; Mars *et al*., 2017). Several theories suggest that the evolutionary path leading to language and tool use in humans may be built upon modifications of circuits that subserve gestures and imitation (Lewis, 2006). Macaques are thought to emulate the goals and intentions of others, whereas chimpanzees can also imitate certain specific actions, but humans have an even stronger bias for high-fidelity copying of precise sequences of actions, which has been called “overimitation” (Hecht *et al*., 2013). Our findings provide a potential explanation for these phenomena in that the macaques showed no asymmetric network connections, the chimpanzees showed a few asymmetric connections, but the humans showed a large number of asymmetric connections. These species differences in leftward asymmetric connections involving language and tool use may reflect human specializations for language and complex tool use.

Unlike the humans, who showed considerable leftward asymmetry connectivity between the IPL and the lateral temporal cortex, the chimpanzees showed few leftward asymmetric connections between the IPL and the temporal cortex, including the planum temporale, aSTG, and aSTS, while macaques showed symmetric connections between the IPL and temporal cortex. The planum temporale is considered to include part of Wernicke’s area homolog (Spocter *et al*., 2010), and displays leftward anatomical asymmetry in humans and great apes (Gannon *et al*., 1998; Hopkins *et al*., 1998). Recent work suggested a left-hemispheric size predominance of the planum temporale in olive baboons, a nonhominid primate species (Marie *et al*., 2018). We speculate that this planum temporale asymmetry may not be the only prominent characteristic related to language lateralization. The patterns from symmetry in macaques to asymmetry in humans and chimpanzees in the present study provide a possible new evidence that neural connectivity asymmetry may underlie the roots of language specialization, with the initial emergence of hemispheric specializations in apes which are elaborated even further in human brain evolution. In addition, identifying increased asymmetric connections between the IPL and planum temporale in human brains compared to chimpanzees and macaques reinforces the evidence that the evolutionary origin of human language capacities is related to further left hemispheric specialization of neural substrates for auditory processing that are shared with other primates (Balezeau *et al*., 2020). Since the aSTG and aSTS have been implicated in semantic and phonologic processing in humans (Vigneau *et al*., 2006), the leftward asymmetric connections of the IPL with the aSTG and aSTS may be relevant to the evolution of human language processing. Our results suggest an evolutionary trajectory for the connectivity of the IPL with the temporal cortex; that is, the connectivity started as symmetric in macaque monkeys, began to develop asymmetrically in chimpanzees, and finally achieved the greatest degree of asymmetry and is refinement in humans. This sequence may support the emergence of language and language-related functions.

### Species-specific differences in asymmetric connectivity in chimpanzees and humans

Species-specific differences in asymmetric connectivity between the IPL and SPL were found in chimpanzees and humans, with leftward asymmetry in the former and rightward asymmetry in the latter, whereas no asymmetry of this connectivity was found in macaques. These species differences in hemispheric asymmetry may reflect evolutionary changes responsible for adaptations or the production of new abilities in the human brain. Structurally, in chimpanzees, right anatomical asymmetry in the white matter below the SPL (Hopkins *et al*., 2010) may increase the right connectivity between the IPL and SPL compared with the left side. In humans, the leftward volumetric asymmetry in the SPL (Goldberg *et al*., 2013), together with leftward volumetric asymmetry in the IPL (the C2), may support the leftward asymmetric connectivity. Functionally, interaction between the IPL and SPL is crucial for tool use, which is dominant in the left hemisphere, and visuospatial function, which is dominant in in the right (Lewis, 2006; De Schotten *et al*., 2011b; Catani *et al*., 2017). As for tool use, in contrast to the relatively simple tools used by chimpanzees and other species, humans can create complex artifacts through a sequence of actions that may incorporate multiple parts, reflecting a deep understanding of the kinematics of our bodies, the mechanical properties of surrounding objects, and the unique demands of the external environments in which we live (Povinelli *et al*., 2000; Johnson-Frey, 2004). In addition, complex tool use requires the SPL to code the location of the limbs relative to other body parts during planning and executing tool-use movements or hand gestures (Wolpert *et al*., 1998; Johnson-Frey *et al*., 2005; Lewis, 2006). Leftward asymmetric connectivity between the IPL and SPL may have provided a connectional substrate for complex tool use during human evolution. As for visuospatial functions, the rightward asymmetric connectivity between the IPL and SPL in chimpanzees may indicate that visuospatial functions are dominant in the right hemisphere and had already been lateralized to the right hemisphere from the common ancestor with macaques. During evolution, these lateralized functions may be retained in the human brain. Meanwhile, the lateralized directional reversal of this connectivity from the right to the left hemisphere may reflect evolutionary adaptations for the emergence of new abilities, such as sophisticated and complex tool making and use.

### Human unique asymmetric connectivity of IPL subregions

Unlike the chimpanzees and macaques, humans showed leftward asymmetry in the connection between the rostral IPL (the C1 and C2) and the primary motor cortex, which is consistent with a larger neuropil volume in the left primary motor cortex than in the right side (Amunts *et al*., 1996). Meanwhile, the leftward asymmetric volume of the anterior IPL and the primary motor cortex may also increase the neural connectivity between these two regions in the left hemisphere compared with the right side. Such a leftward connection is thought to be related to handedness and hand manual skills (Amunts *et al*., 1996; Amunts *et al*., 1997). In contrast to humans, chimpanzees and macaques did not show any asymmetric connectivity between the IPL and the primary motor cortex. A more recent study reported that, in olive baboons, contralateral hemispheric sulcus depth asymmetry of the central sulcus related to the motor hand area is correlated with the direction and degree of hand preference, as measured by a bimanual coordinated tube task, but only about 41% of them were classified as right-handed and 33% were classified as left-handed (Margiotoudi *et al*., 2019). Although previous studies have shown that chimpanzees exhibit population-level handedness in the use of tools and a corresponding asymmetry in the primary motor cortex, inferior frontal cortex, and parietal operculum (Gilissen and Hopkins, 2013; Hopkins *et al*., 2017), they do not show handedness as a more universal trait or exhibit manual dexterity to the same extent as humans. One possible explanation is that humans developed the asymmetric connectivity that became the structural basis for specific behaviors of handedness and hand skills during evolution.

An unexpected finding was that in humans the IPL, particularly the C3, showed rightward asymmetric connectivity with the IFG. Since the IPL and the IFG are interconnected through the SLF3, which is strongly rightward asymmetric (De Schotten *et al*., 2011b), it may also increase the connection between the IPL and IFG in the right hemisphere. Functionally, the left IFG is involved in various aspects of language functions, including speech production and semantic, syntactic, and phonological processing (Wang *et al*., 2020), whereas the right IFG is associated with various cognitive functions, including attention, motor inhibition, and social cognitive processes (Hartwigsen *et al*., 2019). Our result of rightward asymmetry in this connectivity seems to be associated with attention and social function, but not language, although language dominance in the left hemisphere is considered to be a common characteristic in humans.

The widespread asymmetric connections of the IPL in humans compared with the other two primates is in keeping with the inter-hemispheric independence hypothesis, in which, during evolution, brain size expansion led to hemispheric specialization due to time delays in neuron signaling over increasing distances, resulting in decreased inter-hemispheric connectivity and increased intra-hemispheric connectivity (Ringo *et al*., 1994; Phillips *et al*., 2015). While having more cortical neurons (local characteristics) in one hemisphere than the other seems to be a necessary condition for asymmetries of complex and flexible behaviors, it is not a full condition for such behaviors. Given that a function or behavior in an area is determined by its connectivity or networks in which it is involved (Passingham *et al*., 2002), the widespread lateralized connections may provide the human brain with the increased computational capacity necessary for processing language and complex tool use and may play a facilitating role in human cognitive and behavioral specialization.

### Methodological considerations

The three levels of analyses, i.e., the vertex-wise, ROI-wise, and tract-wise analyses, were performed to provide a full description of the connectivity asymmetry. However, it should be noted that some analyses produced results that were not completely consistent with each other. In humans, the connectivity of the IPL with SLF3 and AF was left-lateralized in the C2 while right-lateralized in the C3. The previous studies assessed the asymmetry of the SLF3 and AF with local characteristics such as cortical volume, voxel count, and FA and their average across all the voxels in the tracts (de Schotten *et al*., 2011a; De Schotten *et al*., 2011b; Kamali *et al*., 2014), the SLF3 and AF were usually found to have a single pattern, e.g., leftward asymmetry, rightward asymmetry, or symmetry. However, our results seem to indicate two different asymmetric patterns for the SLF3 and AF, both of which connect the IPL subregion C2 and C3 to the IFG (De Schotten *et al*., 2011b; Hecht *et al*., 2015; Barbeau *et al*., 2020). Furthermore, these connectivity asymmetries matched well with the ROI-wise and tract-wise analyses. The leftward connectivity asymmetry of the human C2 with the SLF3 and AF using the tract-wise approach corresponds to that of the human C2 with the IFG and precentral gyrus using the ROI-wise approach. The rightward connectivity asymmetry of the human C3 with the SLF3 and AF using the tract-wise approach corresponds to that of the human C3 with the IFG using the ROI-wise approach. The SLF3, located at the ventrolateral SLF, connects to the IPL, especially the anterior part, and from there predominantly to the ventral premotor and prefrontal areas (De Schotten *et al*., 2011b; Kamali *et al*., 2014; Barbeau *et al*., 2020). The C2 and C3 appear to separate the SLF3 into two finer components, one connecting the posterior IFG and anterior IPL with leftward asymmetry, and one connecting the anterior IFG to the posterior IPL with rightward asymmetry. The two types of connectivity patterns are consistent with previous studies using invasive tract-tracing findings in macaque monkeys and resting-state functional connectivity results in humans to study frontal and parietal connectivity (Petrides and Pandya, 2009; Margulies and Petrides, 2013). Our results indicated that both cortical areas, such as the IFG, and subcortical tracts, such as the SLF3 and AF, have at least two distinct subcomponents.

The inconsistency was observed when significant ROI-wise connectivity asymmetry was found but few or no significant tract-wise connectivity asymmetries were found. For example, the human C1 showed ROI-wise connectivity asymmetry with the precentral gyrus and insula but no significant tract-wise connectivity asymmetry. The IPL is connected to the precentral gyrus mainly through the SLF3, which in turn is connected to not only the precentral gyrus but also the IFG and MFG in the prefrontal cortex (De Schotten *et al*., 2011b; Kamali *et al*., 2014; Hecht *et al*., 2015; Barbeau *et al*., 2020). The connectivity between the C1 and the precentral gyrus may include only a portion of the SLF3; this may have diluted the laterality effect from the SLF3 because it may include other pathways that were not in our selected tracts and could, thus, have affected the observed lateralization. In other words, the traditionally defined major fiber tracts are not a single bundle but, instead, contain many subcomponents. Therefore, the patterns of lateralization might not yet have been fully explored in our study. A recent work also suggested that the SLF2, SLF3, and AF could be separated into several branches based on their projections into the prefrontal and/or temporal areas (Barbeau *et al*., 2020). This may be true for the other major fiber tracts, such as the ILF (Latini *et al*., 2017), uncinate fasciculus (Hau *et al*., 2017), and cingulum bundle (Jones *et al*., 2013). On the other hand, brain regions, such as the IFG were connected to many fiber tracts, including the SLFII, SLFIII, and AF. Hence, we did not find tract-wise connectivity asymmetry that corresponded to the ROI-wise connectivity asymmetry in the human C1. This was also the case for the chimpanzee and human C4. The creation of a finer tract atlas should be a priority for future work because this would help to map the tract-wise connectivity asymmetry at a higher resolution.

In conclusion, we identified similar topographical maps of the IPL to study structural and connectional asymmetry in macaques, chimpanzees, and humans. We found that the structural asymmetry of the IPL was independent of the allometric scaling of this region. The connectional analysis revealed that humans had the largest connectional asymmetries of IPL subregions compared to macaques and chimpanzees. The regions showing larger asymmetric connections with the human IPL were associated with language, complex tool use, and handedness, which provided potential anatomical substrates for functional and behavioral lateralization in humans. The opposite asymmetric connection between the IPL and SPL in chimpanzees and humans may reflect distinct species-specific modifications to cortical circuits during the course of ape and human evolution.

## Materials and methods

### Human data

Data from 40 right-handed healthy adults (age: 22–35, 18 males) were randomly selected from the S500 subjects release of the Human Connectome Project (HCP) database (Van Essen *et al*., 2013) (http://www.humanconnectome.org/study/hcp-young-adult/). T1-weighted (T1w) MPRAGE images (resolution: 0.7mm isotropic, slices: 256; field of view: 224 × 320; flip angle: 8°), and diffusion-weighted images (DWI) (resolution: 1.25mm isotropic; slices: 111; field of view: 210 × 180; flip angle: 78°; b-values: 1000, 2000, and 3000 s/mm^2^) were collected on a 3 T Skyra scanner (Siemens, Erlangen, Germany) using a 32-channel head coil.

### Chimpanzee data

Data from 27 adult chimpanzees (*Pan troglodytes*, 14 males) were made available by the National Chimpanzee Brain Resource (http://www.chimpanzeebrain.org, supported by the NIH National Institute of Neurological Disorders and Stroke). Data, including T1w and DWI, were acquired at the Yerkes National Primate Research Center (YNPC) on a 3T MRI scanner under propofol anesthesia (10 mg/kg/h) using previously described procedures (Chen *et al*., 2013). All procedures were carried out in accordance with protocols approved by YNPRC and the Emory University Institutional Animal Care and Use Committee (Approval no. YER-2001206).

DWI were acquired using a single-shot spin-echo echo-planar sequence for each of 60 diffusion directions (b = 1000 s/mm^2^, repetition time 5900 ms; echo time 86 ms; 41 slices; 1.8 mm isotropic resolution). DWI with phase-encoding directions (left–right) of opposite polarity were acquired to correct for susceptibility distortion. For each repeat of a set of DWI, five b = 0 s/mm^2^ images were also acquired with matching imaging parameters. T1w images were also acquired for each subject (218 slices, resolution: 0.7×0.7×1mm).

### Macaque data

Data from 8 male adult rhesus macaque monkeys (*Macaca mulatta*) were obtained from TheVirtualBrain (Shen *et al*., 2019). All surgical and experimental procedures were approved by the Animal Use Subcommittee of the University of Western Ontario Council on Animal Care (AUP no. 2008-125) and followed the Canadian Council of Animal Care guidelines. Surgical preparation and anesthesia as well as imaging acquisition protocols have been previously described (Shen *et al*., 2019). Images were acquired using a 7-T Siemens MAGNETOM head scanner. Two diffusion-weighted scans were acquired for each animal, with each scan having opposite phase encoding in the superior-inferior direction at 1 mm isotropic resolution, allowing for correction of susceptibility-related distortion. For five animals, the data were acquired with 2D EPI diffusion, while for the remaining three animals, a multiband EPI diffusion sequence was used. In all cases, data were acquired with b = 1000 s/mm^2^, 64 directions, 24 slices. Finally, a 3D T1w image was also collected for each animal (128 slices, resolution: 0.5 mm isotropic).

### Image preprocessing

The human T1w structural data had been preprocessed following the HCP’s minimal preprocessing pipeline (Glasser *et al*., 2013), while the chimpanzee and monkey T1w structural data had been preprocessed following the HCP’s nonhuman preprocessing pipelines described in previous studies (Glasser *et al*., 2013; Donahue *et al*., 2018). Briefly, the processing pipeline included imaging alignment to standard volume space using FSL, automatic anatomical surface reconstruction using FreeSurfer, and registration to a group average surface template space using the Multimodal Surface Matching (MSM) algorithm (Robinson *et al*., 2014). Human volume data were registered to Montreal Neurological Institute (MNI) standard space and surface data were transformed into surface template space (fs_LR). Chimpanzee volume and surface data were registered to the Yerkes29 chimpanzee template (Donahue *et al*., 2018). Macaque volume and surface data were registered to the Yerkes19 macaque template (Donahue *et al*., 2018).

Preprocessing of the diffusion-weighted images was performed in a similar way in the human, chimpanzee, and macaque datasets using FSL. FSL’s DTIFIT was used to fit a diffusion tensor model for each of the three datasets. Following preprocessing, voxel-wise estimates of the fiber orientation distribution were calculated using Bedpostx, allowing for three fiber orientations for the human dataset and two fiber orientations for the chimpanzee and macaque datasets due to the b-value in the diffusion data.

### Definition of the IPL

The IPL, located at the lateral surface of the ventral posterior parietal lobe, is surrounded by several sulci including the Sylvian fissure, superior temporal sulcus (STS), and intraparietal sulcus (IPS) (von Economo and Koskinas, 1925; Von Bonin, 1947; Bailey *et al*., 1950; Pandya and Seltzer, 1982). In the absence of detailed homologous definitions, it is necessary to use cytoarchitectonic delineations and macroscopic boundaries, such as gyri and sulci, that can be reliably identified in all species as the boundaries of the IPL. The region of interest (ROI) of the IPL was manually drawn on the standard surface template using Connectome Workbench (Glasser *et al*., 2013). In the present study, we restricted the ROI to the lateral surface of the IPL and excluded the cortex buried in the sulci, especially the lateral bank of the IPS and the upper bank of the Sylvian fissure. Rostrally, the IPL borders the vertical line between the Sylvian fissure and the rostral lip of the IPS. Dorsally, the IPL borders the lateral bank of the IPS. Ventrally, the anterior ventral IPL borders the upper bank of the Sylvian fissure. The border of the posterior and ventral IPL is formed by the extension of the Sylvian fissure to the top end of the STS in chimpanzees and macaques but by the extension of the Sylvian fissure to the posterior end of the IPS in humans.

### Connectivity-based parcellation

We used a data-driven connectivity-based parcellation framework modified from Fan *et al* (2016) (**Figure 1**). All steps in the framework were processed on surface data because the surface-based method has advantages, such as cortical areal localization (Coalson *et al*., 2018), over the traditional approach and because the use of surface meshes is a straightforward way to improve existing tractography processing pipelines, such as the precise locations of streamline seeding and termination (St-Onge *et al*., 2018). The surface ROI was first registered to native surface using MSM (Robinson *et al*., 2014). The probabilistic tractography was performed on the native mesh representing the gray/white matter interface using Probtrackx. The pial surfaces were used as stop masks to prevent streamlines from crossing sulci. 5000 streamlines were seeded from each of the white matter surface vertices in the seed region to estimate its whole-brain connectivity profile and were downsampled to 5 mm isotropic voxels to construct the native connectivity M-by-N, a matrix between all the IPL vertices (M) and the brain voxels (N). Based on the native connectivity matrix, a symmetric cross-correlation M-by-M matrix was calculated to quantify the similarity between the connectivity profiles of each IPL vertex. A group cross-correlation matrix was calculated by averaging the cross-correlation matrix across subjects.

Data-driven spectral clustering was applied to the group cross-correlation matrix to define the anatomical boundaries of the IPL. Spectral clustering can capture clusters that have complicated shapes, making them suitable for parcellating the structure of complicated brain regions such as the IPL. In addition, the spectral clustering algorithm was successfully used to establish the Brainnetome Atlas (Fan *et al*., 2016). However, the number of clusters must be defined by the experimenter when using this method. In the current study, we explored from two to twelve parcellations.

### Volumetric analysis of the IPL

The cortical gray matter volumetric measurements were calculated using Freesurfer. Total cortical volumes were determined by the space between the white and pial surfaces in native space. Each subregion drawn on standard surface space was registered to native surface space using an existing mapping between the two meshes. The volume of the IPL and its subregions was determined by averaging all the vertices for each subject.

### Functional decoding of each subregion of the human IPL

Each subregion was first mapped to MNI volume space using a ribbon-constrained method in Connectome Workbench. To decode the functions of each subregion, we used the automated meta-analysis database, Neurosynth (Yarkoni *et al*., 2011) to identify the terms that were the most associated with each subregion. The top five non-anatomical terms with the highest correlation values were kept for all subregions and redundant terms, such as ‘*semantic*’ and ‘*semantics*’, were only considered once. For simplicity, we only showed the positive correlations found by decoding because negative correlations do not directly inform us about the functions of the subregions. The lateralization for each term was obtained by calculating the difference in the correlation values of the subregions between the left and right hemispheres.

### Mapping anatomical connectivity profiles

To map the whole-brain anatomical connectivity pattern for each cluster, we performed probabilistic tractography by drawing 5000 samples from each vertex in each cluster. The resulting tractograms were log-transformed, normalized by the maximum, and then projected onto surface space using the ‘surf_proj’ command in FSL to obtain tractograms in surface space. The surface tractograms were smoothed using a 4 mm kernel for humans, 3 mm kernel for chimpanzees, and 2 mm kernel for macaques. We subsequently averaged the surface tractograms across subjects for the left and right hemispheres separately to obtain population tractograms, which were thresholded by a value of 0.5 for humans, 0.2 for chimpanzees, and 0.3 for macaques due to data quality. The resultant population tractograms represented approximately twenty percent of the non-zero vertexes in the non-thresholded population tractograms and were used for the vertex-wise and ROI-wise comparisons. The volumetric tractograms were used for the tract-wise comparison.

### Vertex-wise analysis

For each subregion, we restricted the analysis to the group mask defined by the combination of the left and mirrored right population tractograms described above. We here used the connectivity probabilistic value to quantify the connectivity between the IPL and each vertex of the rest of the brain. A higher value in the vertex means a higher likelihood of being connected to the IPL than other vertices.

### ROI-wise analysis

Although previous studies have devoted much effort to establishing homologous regions in primates, these are still limited to a few regions, particularly in chimpanzees. To make comparisons across species possible, here we used the common principle of macroscopic anatomical boundaries based on the gyri and sulci to define ROIs in the cerebral cortex. Specifically, the Desikan–Killiany–Tourville (DKT) atlas was used for humans (Desikan *et al*., 2006), a modified DKT atlas for the chimpanzees, and the Neuromaps atlas for the macaques (Rohlfing *et al*., 2012). Because the Neuromaps atlas is volumetric, we first mapped it to surface space for the subsequent calculations. A total of eleven cortical ROIs were chosen for each hemisphere: the superior frontal gyrus (SFG), inferior frontal gyrus (IFG, a combination of the pars triangularis and pars opercularis in humans and chimpanzees), anterior cingulate gyrus (CGa, a combination of the rostral and caudal anterior-cingulate in humans and chimpanzees), orbitofrontal cortex (Orb), precentral gyrus (PreG), postcentral gyrus (PostG), superior parietal lobule (SPL), precuneus, superior temporal gyrus (STG), middle temporal gyrus and inferior temporal gyrus (MTG/ITG), and insula. The MTG/ITG was a combination of the MTG and ITG in humans and chimpanzees due to the absence of the MTG in macaques. The connectional value for each ROI was calculated by averaging all vertices in the ROI on the individual surface tractogram for each subregion.

### Tract-wise analysis

To investigate which subcortical fiber tracts are associated with lateralization of cortical areas connected to the IPL, we analyzed the lateralization of the subcortical white matter tracts connected to the IPL across species. A total of seven tracts were chosen: the three branches of the superior longitudinal fasciculus, arcuate fasciculus, middle longitudinal fasciculus, inferior longitudinal fasciculus, inferior fronto-occipital fasciculus. The automated tractographic protocols for tracts for each species were from previous studies (Bryant *et al*., 2020) and these tracts were reconstructed using the Xtract tool (Warrington *et al*., 2020). The mean value for each tract was calculated by averaging all voxels in the tract in the individual volumetric tractogram for each subregion.

### Statistical analysis

To investigate the allometric relationship between the volume of each of the IPL subregions and the total gray matter volume using log-transformed data (Donahue *et al*., 2018), linear regression was performed by pooling the human, chimpanzee, and macaque data for each of the IPL subregions, separately. To test whether the scaling regression slopes differed significantly between the two hemispheres, we performed an ANCOVA for comparisons across the two regression slopes for each plot.

In all the analyses of the structural and connectional asymmetries (i.e., volumetric, vertex-wise, ROI-wise, and tract-wise), the asymmetry index (AI) was defined as the difference between values for the left and right hemispheres according to the formula AI = 2 x (R − L) / (R + L). For the vertex-wise analysis, a one-sample *t* test was performed at each vertex on the group mask for each species using PALM, with 5000 permutations with a sign-flip strategy (Winkler *et al*., 2014). The statistically significant level was set at false discovery rate corrected *p* < .05. The effect sizes (Cohen’s *d*) were displayed on the average surface. For the volumetric, ROI-wise, and tract-wise analysis, a two-sided Wilcoxon signed-rank test was performed for each subregion. Bonferroni correction was then used for multiple comparisons for seeds, ROIs or tracts, and species, with statistical significance set at *p* < .05.

## Acknowledgments

This work was partially supported by the National Natural Science Foundation of China (Grant Nos. 91432302, 82072099, and 31620103905), the Science Frontier Program of the Chinese Academy of Sciences (Grant No. QYZDJ-SSW-SMC019), Beijing Municipal Science & Technology Commission (Grant Nos. Z161100000216139 and Z171100000117002), the Guangdong Pearl River Talents Plan (2016ZT06S220), Key-Area Research and Development Program of Guangdong Province (2018B030333001), the Youth Innovation Promotion Association, the Beijing Advanced Discipline Fund, and the National Science Foundation (SMA-1542848). The National Chimpanzee Brain Resource was supported by NIH - National Institute of Neurological Disorders and Stroke (NIH Grant No. NS092988). We thank Rhoda E. and Edmund F. Perozzi, PhDs, for English language and editing assistance.

## Author contributions

T.J. and L.F. designed the study. L.C. and G.L. analyzed the data. C.C.S collected the chimpanzee data. L.C. wrote the first draft of the manuscript. T.J. supervised the study. All authors revised and approved the manuscript.

## Data availability

The datasets analyzed during the current study are available at https://www.humanconnectome.org, http://www.chimpanzeebrain.org, http://openneuro.org/datasets/ds001875/versions/1.0.3, and http://www.neurosynth.org.

