## Supplementary for "Connectional Asymmetry of the Inferior Parietal Lobule Shapes Hemispheric Specialization in Humans, Chimpanzees, and Rhesus Macaques"

### SUPPLEMENTARY INFORMATION

#### Supplementary results

##### Functional decoding of human IPL subregions

To decode the functions of the human IPL subregions, we investigated the relationship of each subregion with brain activation patterns obtained from the Neurosynth database (Yarkoni et al., 2011). The functional decoding of each subregion in each hemisphere is shown in **Supplementary Figure 2**. To assess in which hemisphere a given functions was dominant, we calculated the differences in the correlation values for the left and right subregions (**Supplementary Figure 2C**). The five most correlated terms for the left and right corresponding subregions were shown to be similar in some cases and dissimilar in others, suggesting functional lateralization in each subregion. These selected items exhibited a dichotomous pattern of asymmetry. Specifically, the left and right C1 both demonstrated relatively high correlations with sensory-related terms, such as *somatosensory*, *tactile*, *touch*, and *pain*. The term *tool* showed a prominently higher correlation with the left C1 compared to the right C1. The left C2 demonstrated relatively high correlations with terms including *grasping*, *monitoring*, *inhibition*, *semantics*, and *tool*; and the right C2 showed relatively high correlations with terms including *nogo*, *inhibition*, *preparatory*, *error*, and *detection*. The term *tool* and the language-related term *semantics* showed relatively high correlations with the left C2, whereas executive-related terms, such as *nogo* and *inhibition*, showed relatively high correlations with the right C2. The left C3 demonstrated relatively high correlations with terms including *retrieval*, *solving*, *recollection*, *judgments*, and *coherent*; and the right C3 showed relatively high correlations with terms including *beliefs*, *intentions*, *mentalizing*, *monitoring*, and *perspective*. The memory- and language-related terms, such as *retrieval*, *episodic*, *recollection*, *memories*, and *coherent*, showed relatively high correlations with the left C3, whereas executive-related terms, such as *nogo* and *inhibition*, and the social-related term *beliefs*, were correlated with the right C3. The left C4 demonstrated relatively high correlations with terms including *episodic*, *autobiographical*, *retrieval*, *coherent*, and *memories*; and the right C4 showed relatively high correlations with terms including *spatial*, *attention*, *mentalizing*, *retrieval*, and *relevance*. The memory-related term *episodic* and the language-related term *coherent* showed relatively high correlations with the left C4, whereas the attention- and social-related terms, such as *spatial*, *attention*, *mentalizing*, and *relevance*, were correlated with the right C4.

**Connectivity profiles of IPL subregions**

The whole-brain connectivity profiles of the left and right subregions were mapped in macaques, chimpanzees, and humans using probabilistic tracking (**Supplementary Figure 3**). In macaques, all regions showed consistent connections with the ventral bank of the principal sulcus, the medial frontal cortex, superior parietal lobule (SPL), precuneus, superior temporal sulcus (STS), and insula. The C1, C2, and C3 were also connected with the premotor and sensorimotor cortex. The caudal subregions (the C3 and C4) were more connected with the superior temporal gyrus (STG). Visualization of the connectivity patterns did not show obvious interhemispheric asymmetry. In chimpanzees, all regions were connected with the middle frontal gyrus (MFG), inferior frontal gyrus (IFG), SPL, precuneus, planum temporale, and insula. The rostral subregions (the C1 and C2) were connected with the ventral part of the premotor and sensorimotor cortex, while caudal subregions (the C3 and C4) had strong connections with almost all the STG, the middle temporal gyrus (MTG), and the occipitotemporal areas. Visualization of the connectivity patterns did not show interhemispheric asymmetry. In humans, all regions showed connections with the IFG, the caudal part of the MFG, SPL, dorsal precuneus, and the lateral temporal cortex. The most rostral subregion C1 also showed connections with the primary motor and sensorimotor cortices and relatively few connections with the lateral temporal cortex compared with the other three subregions. The caudal subregion C4 also showed connections extending to the occipitotemporal areas. Visual inspection of the connectivity profiles of the IPL showed obvious interhemispheric asymmetries, especially in connections with the IFG and lateral temporal cortex.

### Supplementary Figures

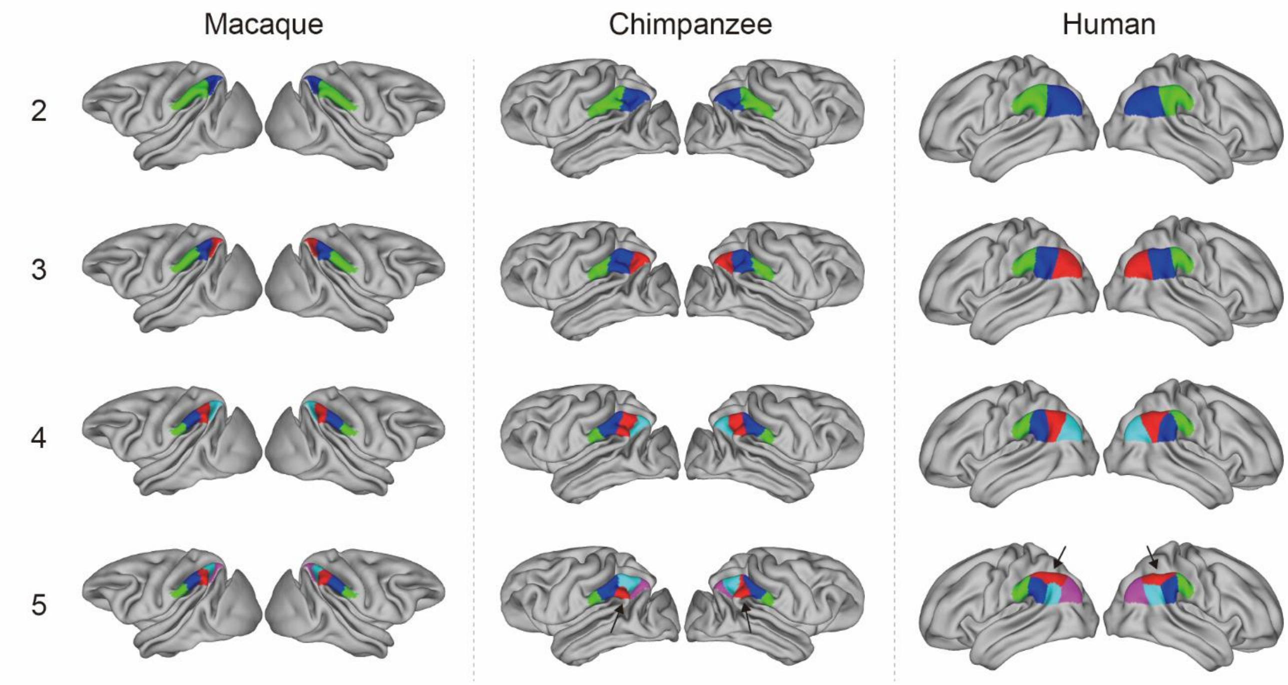


**Supplementary Figure 1.** Two- to five-cluster parcellation of the IPL. The two- to four-cluster solutions showed a consistent rostral-caudal pattern. In the five-cluster solution, a ventral cluster emerged in chimpanzees and a dorsal cluster emerged in humans.

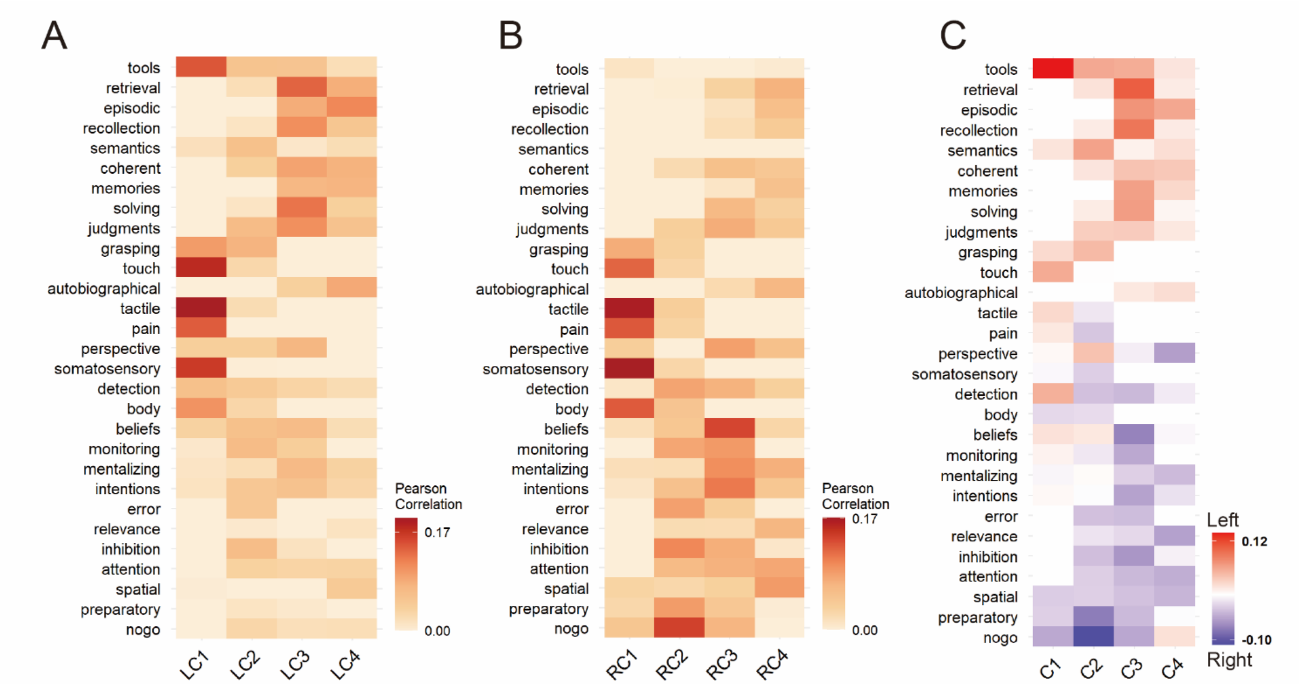


**Supplementary Figure 2.** Functional decoding of the human left (**A**) and right (**B**) inferior parietal lobule (IPL) subregions. (**C**) Differences between the correlation values of the left and right IPL subregions.

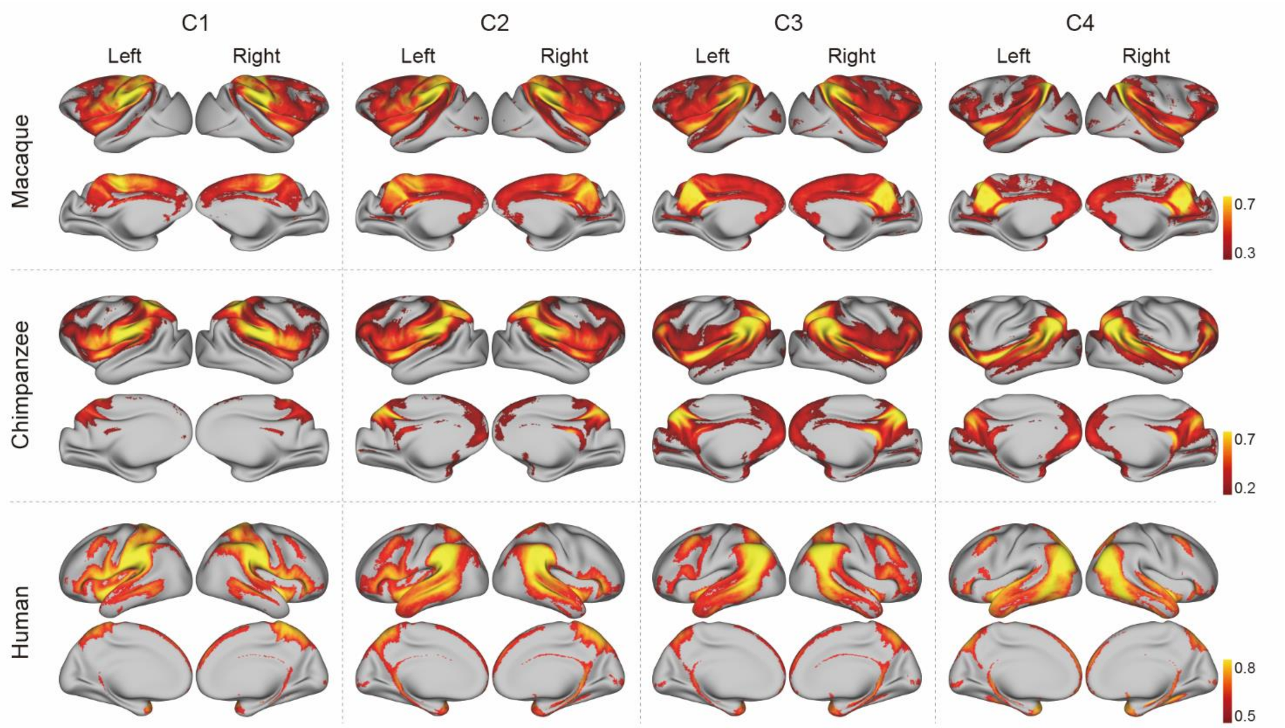


**Supplementary Figure 3.** Connectivity profiles of IPL subdivisions across species. Probabilistic tractography was performed for each IPL subdivision to map its whole-brain connectivity profiles. The group tractograms shown were at a threshold of 0.3 for macaques, 0.2 for chimpanzees, and 0.5 for humans.

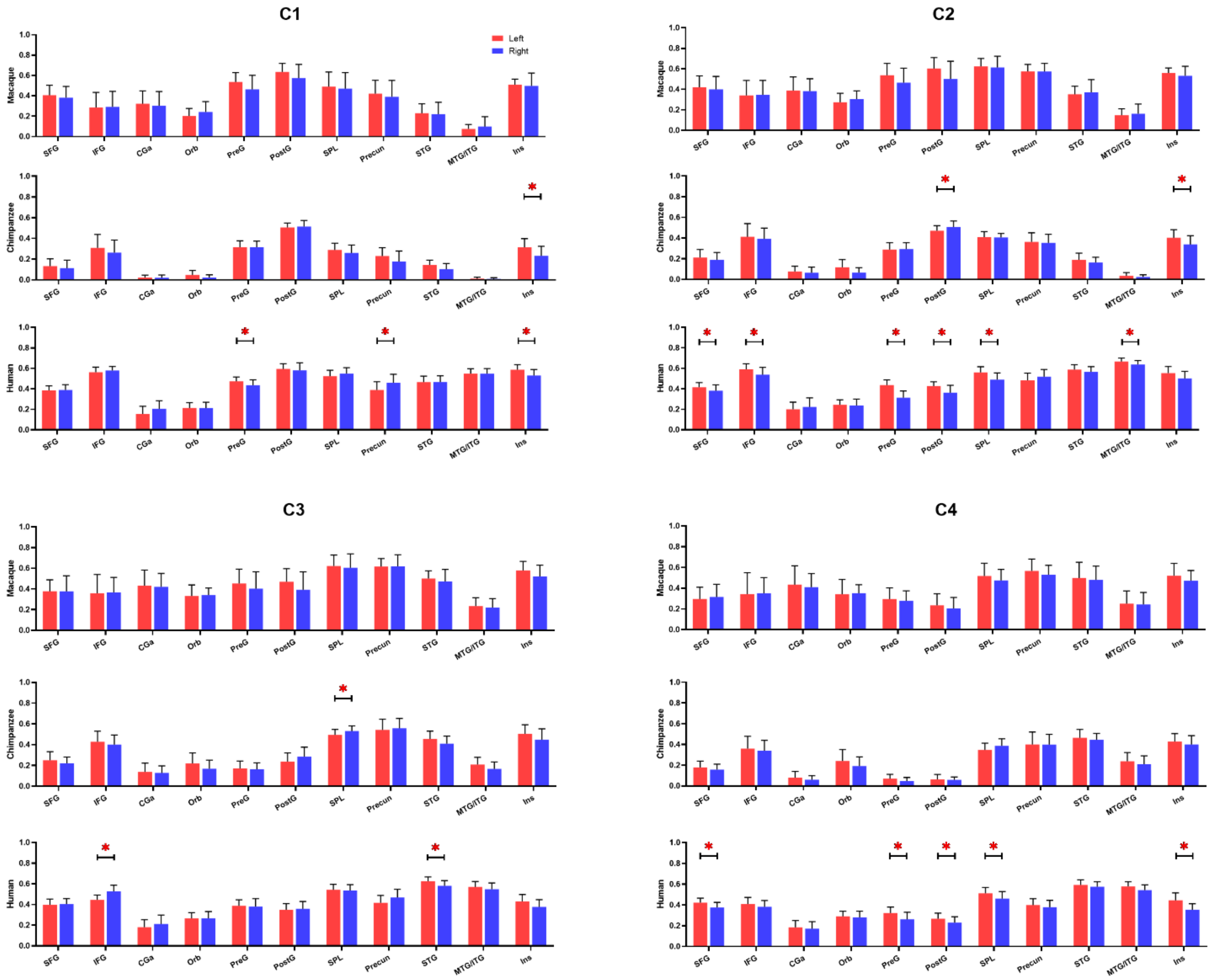


**Supplementary Figure 4.** Bar graphs of the average connectivity values between the inferior parietal lobule (IPL) subregions and 11 cortical regions for each species. The error bars indicate standard deviation. * indicates significance at a Bonferroni corrected level of $p < .05$.

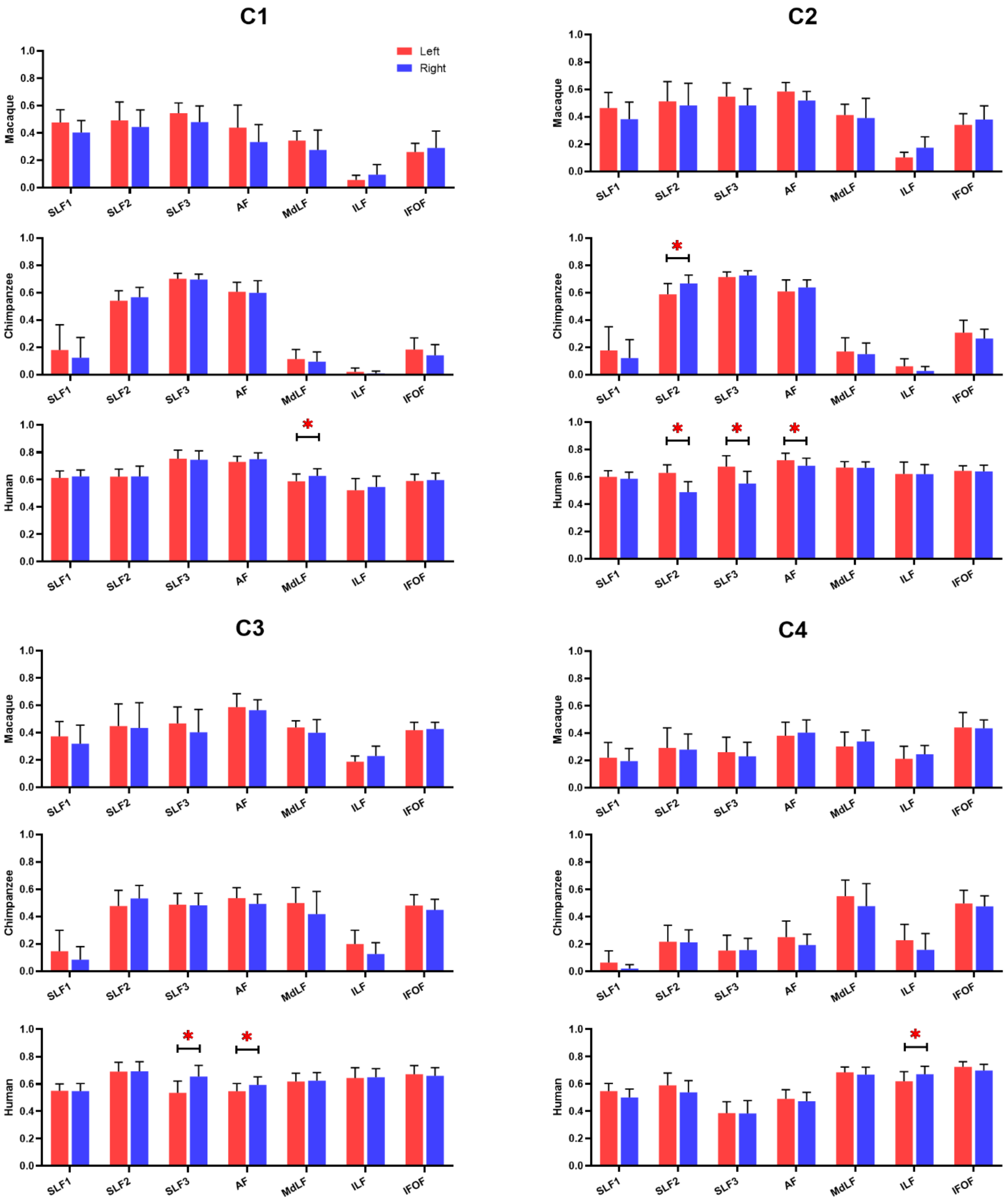


**Supplementary Figure 5.** Bar graphs of the average connectivity values between the inferior parietal lobule (IPL) subregions and 11 subcortical tracts for each species. The error bars indicate standard deviation. * indicates significance at a Bonferroni corrected level of $p < .05$.
